# The effects of the angiotensin II receptor antagonist losartan on appetitive versus aversive learning

**DOI:** 10.1101/472050

**Authors:** Erdem Pulcu, Lorika Shkreli, Carolina Guzman Holst, Marcella L. Woud, Michelle G. Craske, Michael Browning, Andrea Reinecke

## Abstract

Exposure therapy is a first-line treatment for anxiety disorders but remains ineffective in a large proportion of patients. A proposed mechanism of exposure involves a form of inhibitory learning where the association between a stimulus and an aversive outcome is suppressed by a new association with an appetitive or neutral outcome. The blood pressure medication losartan augments fear extinction in rodents and might have similar synergistic effects on human exposure therapy, but the exact cognitive mechanisms underlying these effects remain unknown. In this study, we used a reinforcement learning paradigm with compound rewards and punishments to test the prediction that losartan augments learning from appetitive relative to aversive outcomes. Healthy volunteers (N=53) were randomly assigned to single-dose losartan (50mg) versus placebo. Participants then performed a reinforcement learning task which simultaneously probes appetitive and aversive learning. Participant choice behaviour was analysed using both a standard reinforcement learning model and by analysis of choice switching behaviour. Losartan significantly reduced learning rates from aversive events (losses) when participants were first exposed to the novel task environment, while preserving learning from positive outcomes. The same effect was seen in choice switching behaviour. Losartan enhances learning from positive relative to negative events. This effect may represent a computationally defined neurocognitive mechanism by which the drug could enhance the effect of exposure in clinical populations.

## Introduction

Exposure therapy, often thought of as an analogue to fear extinction, is the first-line intervention for anxiety disorders (1). However, treatment only leads to recovery in 50-60% of patients (2), with one half relapsing back into anxiety at some later point (3, 4). A proposed mechanism of treatment action is a specific form of inhibitory learning where the previous association of a stimulus with ‘threat’ is overwritten by a new association with ‘safety’ (5-7). Within a reinforcement learning framework, this process can be described by the two learned associations being independently driven by prediction errors, produced by the presence or absence of aversive and appetitive events respectively. Strikingly, anxious individuals show deficits in this type of learning, reflected in elevated fear response to threat *and* safety stimuli during fear extinction (8-11). Such findings may explain why response rates to treatment remain low, and they suggest that identifying strategies that compensate for such impairments may improve treatment response rates (6).

Recent pre-clinical and clinical work suggests that drugs targeting the renin-angiotensin system, such as the antihypertensive drug losartan, may represent a novel approach to the augmentation of exposure therapy. In a rodent model, these drugs have been shown to enhance fear extinction (12), and observational data from a large patient cohort indicates that use of angiotensin-converting enzyme inhibitors or angiotensin receptor blockers such as losartan while experiencing trauma is linked to developing fewer traumatic symptoms in the aftermath of that event (13). From a mechanistic perspective, these effects may be driven by central dopaminergic transmission, as angiotensin and dopamine receptors have an overlapping neural topography, being collocated in brain areas involved in reward processing and fear learning (14-19). Further, angiotensin receptors have been directly implicated in the modulation of reward-related dopamine release in the striatum (20), with the angiotensin II receptor antagonist losartan increasing excitatory dopamine D1 receptor activation (21) and preventing dopaminergic cell deterioration in Parkinson’s disease (22).

These findings suggest a specific mechanism of action for losartan which spans its neural, cognitive and behavioral effects: that it facilitates central dopaminergic transmission leading to enhanced learning from positive relative to negative events, which in turn produces a greater impact of exposure therapy. In this study, we take a computational approach to test the cognitive prediction arising from this proposal in human participants. We used an established reinforcement learning task that probes appetitive and aversive learning processes simultaneously, to investigate the degree to which losartan influenced learning from positive relative to negative outcomes (23, 24). In line with the research reported above, we hypothesised that losartan would modulate the priority of learning between aversive and appetitive outcomes, by increasing reward, relative to loss learning rates.

## Methods and Materials

### Participants

Fifty-three healthy participants were recruited through local advertisements. Due to the lack of previous evidence regarding the effect of losartan on learning outcomes, we estimated sample size based on the only available study on the effect of losartan on cognitive function in healthy volunteers. With observed prospective memory detection rates of M=3.9/SD=2.4 after placebo and M=5.7/SD=1.6 after losartan (25), calculations suggested 21 participants per group to achieve an effect size *d*=0.9 and statistical power of 80% (α-level 0.05). We aimed for group sizes of 26 to account for potential drop-out.

Participants were excluded from participation if they met criteria for a current DSM-IV Axis I disorder as assessed using the Structured Clinical Interview for DSM-IV (26). Participants also had to be medication-free for at least 6 weeks, have a body mass index of 18-30 kg/m^2^, and have no first-degree family member with a history of severe psychiatric disease. The study was approved by the Oxford University research ethics committee. All participants gave written informed consent.

### Materials and Study Design

To characterise the sample, all participants completed a questionnaire battery at screening: the Anxiety Sensitivity Index Revised (ASI-R) (27), the neuroticism subscale of the Eysenck Personality Inventory (EPI) (28), the Behavioural Inhibition Scale (BIS) (29), the Beck Depression Inventory II (BDI) (30), and the Attentional Control Scale (ACS) (31). The National Adult Reading Test (NART) (32) was applied to estimate verbal intelligence. On the test day, participants were stratified for gender and randomly allocated to one of two treatment conditions in a double-blind design: losartan (Cozaar, Merck Sharp & Dohme Ltd.) given as a single oral 50mg dose or matched placebo (microcrystalline cellulose; Rayotabs, Rayonex GmbH). Testing took place 1 hour after medication administration, when drug peak plasma levels are typically reached (33, 34).

To assess potential changes in subjective state mood and physiological symptoms, participants completed visual analogue scales before administration of medication and before testing, at which time we also measured heart rate and blood pressure, using an Omron 705IT sphygmomanometer. At the end of the testing session, both participant and experimenter guessed whether losartan or placebo had been given.

### Information Bias Learning Task (IBLT)

This task has been described previously (23) and was presented on a laptop computer running Psychtoolbox software version 3.0 on MATLAB (2015b; MathWorks Inc.). On each trial, two abstract shapes (Agathodaimon font letters) were presented, and a win outcome (15p) and a loss outcome (−15p) were associated with each shape. Participants were asked to choose the shape which they believed would give the better outcome. The outcomes were independent, meaning that a particular shape could be associated with one, both or neither of the win and loss outcomes (Figure 1C). As the two outcomes were independent, participants had to separately learn which shape the win outcome and which shape the loss outcome was likely to be associated with and integrate this information to judge which option was the better one to select in the current trial.

**Figure 1.**
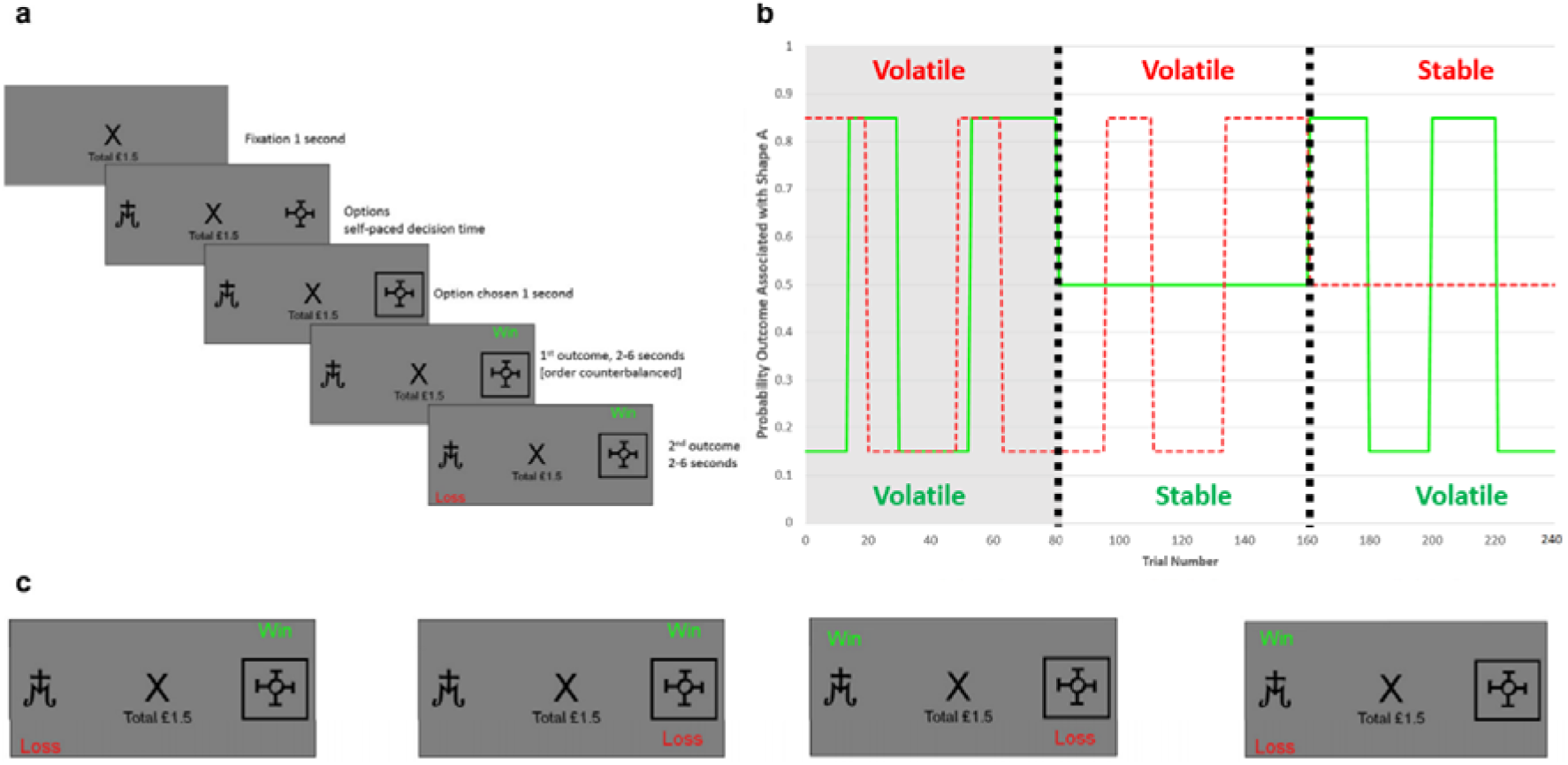
Experimental task design. (A) Timeline of one trial from the learning task. On each trial, participants were presented with two abstract shapes (e.g. shape ‘A’ and ‘B’) and had to choose one. (B) Each shape had win and loss probability which were independent. We manipulated the volatility of these outcome probabilities in different task blocks (black vertical dashed lines separating each block containing 80 trials). In the first block (i.e. the first 80 trials of the task, the light grey area) both outcomes were always equally volatile, which allowed an assessment of participants’ learning bias. In the last two blocks one outcome was volatile while the other was stable. The y-axis represents the probability, p, that an outcome (win in solid green or loss in dashed red) will be found under shape ‘A’ (the probability that it is under shape ‘B’ is 1-p). (C) The independence of win and loss outcome probabilities means that a participant choosing shape A can experience four different outcomes. The independence of win and loss outcomes also meant that the participants had to separately estimate where the win and where the loss would be on each trial in order to perform optimally in this task. This manipulation made it possible to separately estimate learning from the win and loss outcomes. Figure adapted from Pulcu & Browning, 2017.

The task consists of two distinct phases (Figure 1B): an initial block of 80 trials in which both the win and loss outcome reversed frequently (i.e. were volatile) and two further blocks of 80 trials each in which one outcome reversed frequently while the other remained stable. The order of the final two blocks was randomized across participants. The rationale for varying the volatility of the outcomes has been described previously (23). In brief, outcomes which are volatile are more informative than stable outcomes and should lead to participants employing a higher learning rate. Therefore, the first block measures the degree to which participants learn preferentially from positive relative to negative outcomes, whereas the final two blocks estimate how participants adjust their learning to match the volatility of the outcomes. The effect of losartan could manifest either as differential learning in block 1 if it acted to specifically bias learning of affective outcomes, or as an interaction with volatility in the final two blocks if it acted to bias the adaptation of learning to the outcome information content.

The task was self-paced with an explicit rest-session between blocks. The same two stimuli (“Shape A” and “Shape B”) were used within each block but changed between blocks. On average, within each block both outcomes were associated with Shape A on 50% of trials (the probability for Shape B is 1 - Shape A, therefore outcomes were also associated with Shape B on 50% of trials). For the stable block this was achieved with a constant association between the outcome and Shape A of 50%, whereas in the volatile block the association shifted between 15 and 85% every 14 to 30 trials. Choice data from the task were analysed in two ways: by fitting a reinforcement learning model and using a non-model-based approach.

### Model-Based Analysis of Participant Choice Behaviour

In line with our previous work (23), we analysed choice behaviour using a model in which a Rescorla-Wagner (35) learning component was coupled to a soft-max action selector. This model separately estimates the probability of the two outcomes (*rwin* and *rloss*) being associated with Shape A on trial *i*.

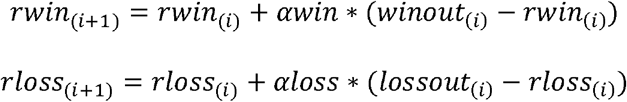

Each process has its own learning rate (*awin* and *aloss*) which may take values between 0 and 1. *winout* and *lossout* represent the outcome of each trial, being 1 if the relevant outcome was associated with Shape A and 0 otherwise. *rwin* and *rloss* were initialised at 0.5. These two estimated probabilities fed into an action selection stage:

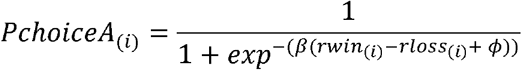

Here, *PchoiceA* is the probability the model will select Shape A on the current trial, *β* is an inverse temperature term which controls the degree to which the model’s choices are driven by the learned probabilities, and *ϕ* is a bias term which allows the model to prefer one shape over the other. In total, this model has 4 free parameters (two learning rates, one inverse temperature and one bias parameter). Parameter values were estimated by calculating the joint posterior probability of parameters over the parameter space, given the choice of participants (23, 24). The estimated value of each parameter was then calculated as the expected value of the parameter’s marginal distribution. Separate parameters were estimated for each participant and each block. Details of the comparison with alternative models are provided in the supplementary materials.

### Non-Model-Based Analysis of Participant Choice Behaviour

This approach can complement model-based analyses by illustrating the behavioural effect of the intervention in the absence of the assumptions associated with specific models (23). The IBLT task includes trials in which both the win and loss outcomes are associated with the same shape (Figure 1C). These trials are particularly useful as the choice made on the next trial provides a measure of the relative impact of the two outcomes on behaviour.

Specifically, if a participant is more influenced by wins than losses, then they would be more likely to stick with the choice they made if it was associated with both outcomes, and to switch if their choice was associated with neither outcome. In contrast, if they were more influenced by losses then they would show the opposite pattern, switching more often after choosing the shape associated with both outcomes. The simplicity of this analysis comes at the cost of precision—unlike model-based approaches it is unable to differentiate an increase in the influence of wins from a decrease in the influence of losses and only reports the relative influence of wins versus losses.

### Statistical Analysis

We used two-tailed tests and □=0.05 in SPSS 25 software (IBM SPSS, Inc., Armonk NY). Potential drug-induced changes in physiological and VAS parameters were assessed using 2 time (baseline, peak) x 2 group (placebo, losartan) mixed model ANOVAs. Choice behaviour from the first block of the IBLT task was analysed separately from the last two blocks as these assess different effects. The effect of losartan on the parameters derived from the model-based analyses were assessed using mixed-model ANOVAs with the within-subject factors valence (positive, negative) and block (only for analysis of last 2 blocks; win volatile, loss volatile) and the between-subject factors group (losartan, placebo) and block order (only for analysis of last 2 blocks; win volatile first or second). A generalized estimating equation framework using a logistic-link function was used to analyse non-model-based results, as the dependent variable is the proportion of trials in which choice switching occurred. Within-subject factors were trial type (both outcomes associated with chosen shape, neither outcomes associated with chosen shape) and block type (for analysis of last 2 blocks only) with between-subject factors group and block order. Reaction times were Box-Cox transformed (36). Learning rates were transformed to the infinite real line before analyses, using inverse logistic transform (results reported in normal space for ease of interpretation).

## Results

### Demographics and Drug Side Effects

The groups (placebo N=25, losartan N=28) were well-balanced on sociodemographic and clinical parameters. There were no differences in heart rate, blood pressure and VAS changes from baseline to drug-peak level, all *F*(1,51)<1.83, all *p*>.18 (Table 1, 2).

**Table 1:**
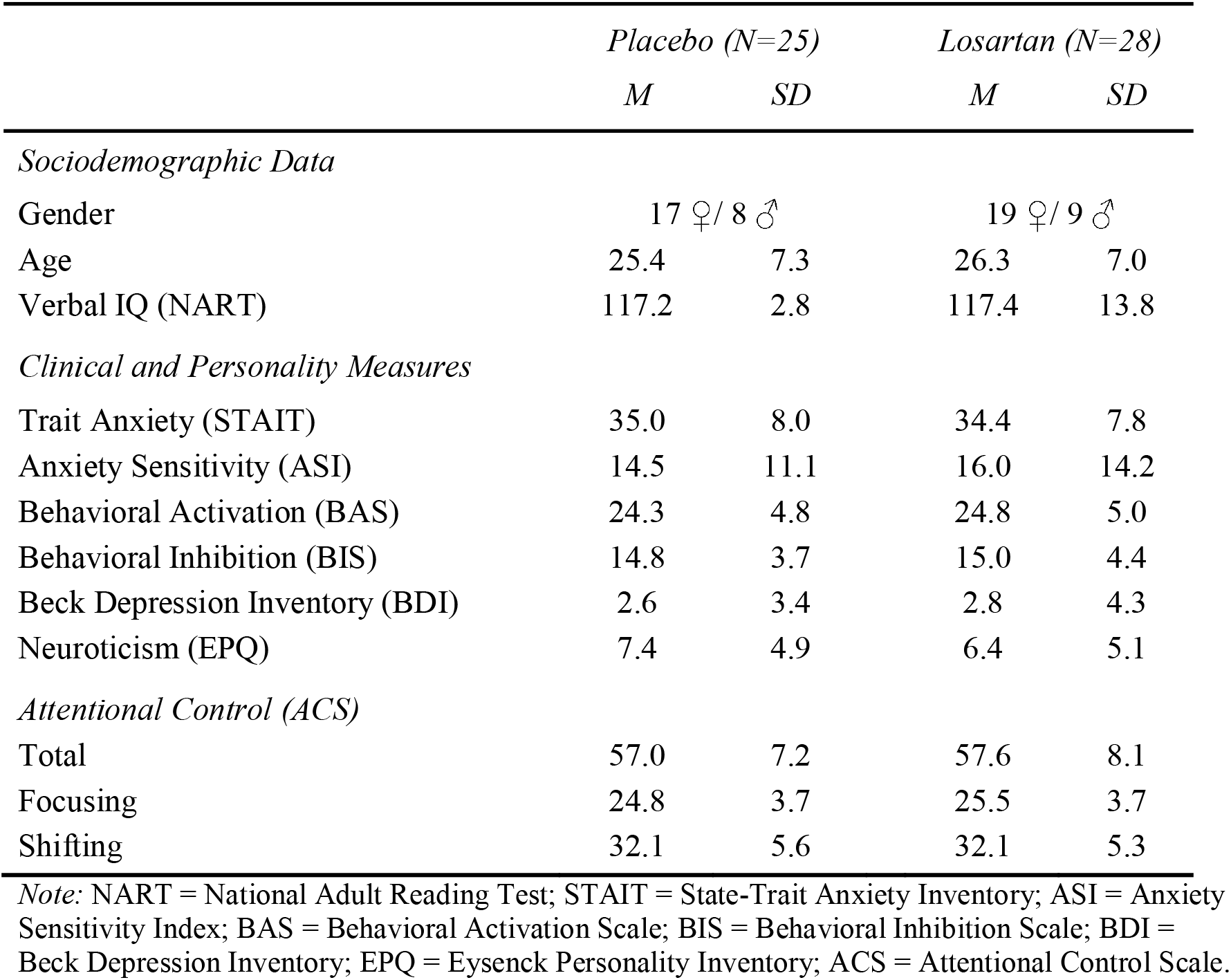
Sociodemographic and clinical characteristics of participants in the losartan versus placebo group (*M, SD).*

|  | *Placebo (N=25)* |  | *Losartan (N=28)* |  |
| --- | --- | --- | --- | --- |
|  | *M* | *SD* | *M* | *SD* |
| *Sociodemographic Data* |  |  |  |  |
| Gender | 17 ♀ / 8 ♂ |  | 19 ♀ / 9 ♂ |  |
| Age | 25.4 | 7.3 | 26.3 | 7.0 |
| Verbal IQ (NART) | 117.2 | 2.8 | 117.4 | 13.8 |
| *Clinical and Personality Measures* |  |  |  |  |
| Trait Anxiety (STAIT) | 35.0 | 8.0 | 34.4 | 7.8 |
| Anxiety Sensitivity (ASI) | 14.5 | 11.1 | 16.0 | 14.2 |
| Behavioral Activation (BAS) | 24.3 | 4.8 | 24.8 | 5.0 |
| Behavioral Inhibition (BIS) | 14.8 | 3.7 | 15.0 | 4.4 |
| Beck Depression Inventory (BDI) | 2.6 | 3.4 | 2.8 | 4.3 |
| Neuroticism (EPQ) | 7.4 | 4.9 | 6.4 | 5.1 |
| *Attentional Control (ACS)* |  |  |  |  |
| Total | 57.0 | 7.2 | 57.6 | 8.1 |
| Focusing | 24.8 | 3.7 | 25.5 | 3.7 |
| Shifting | 32.1 | 5.6 | 32.1 | 5.3 |

*Note:* NART = National Adult Reading Test; STAIT = State-Trait Anxiety Inventory; ASI = Anxiety Sensitivity Index; BAS = Behavioral Activation Scale; BIS = Behavioral Inhibition Scale; BDI = Beck Depression Inventory; EPQ = Eysenck Personality Inventory; ACS = Attentional Control Scale.

**Table 2.**
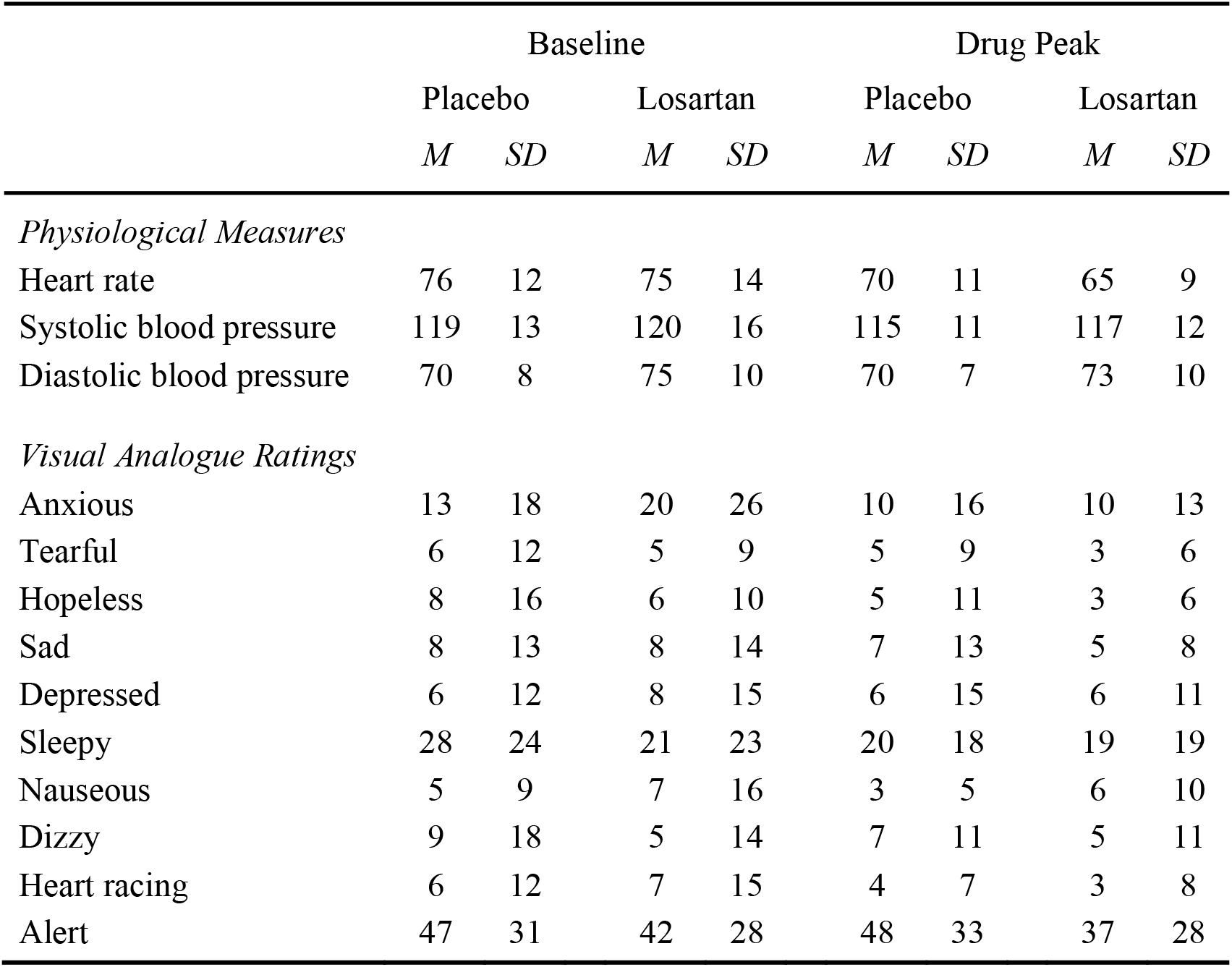
Heart rate, blood pressure and visual analogue scale ratings in the two groups before drug intake and at drug peak-level.

|  | Baseline |  |  |  | Drug Peak |  |  |  |
| --- | --- | --- | --- | --- | --- | --- | --- | --- |
|  | Placebo |  | Losartan |  | Placebo |  | Losartan |  |
|  | *M* | *SD* | *M* | *SD* | *M* | *SD* | *M* | *SD* |
| *Physiological Measures* |  |  |  |  |  |  |  |  |
| Heart rate | 76 | 12 | 75 | 14 | 70 | 11 | 65 | 9 |
| Systolic blood pressure | 119 | 13 | 120 | 16 | 115 | 11 | 117 | 12 |
| Diastolic blood pressure | 70 | 8 | 75 | 10 | 70 | 7 | 73 | 10 |
| *Visual Analogue Ratings* |  |  |  |  |  |  |  |  |
| Anxious | 13 | 18 | 20 | 26 | 10 | 16 | 10 | 13 |
| Tearful | 6 | 12 | 5 | 9 | 5 | 9 | 3 | 6 |
| Hopeless | 8 | 16 | 6 | 10 | 5 | 11 | 3 | 6 |
| Sad | 8 | 13 | 8 | 14 | 7 | 13 | 5 | 8 |
| Depressed | 6 | 12 | 8 | 15 | 6 | 15 | 6 | 11 |
| Sleepy | 28 | 24 | 21 | 23 | 20 | 18 | 19 | 19 |
| Nauseous | 5 | 9 | 7 | 16 | 3 | 5 | 6 | 10 |
| Dizzy | 9 | 18 | 5 | 14 | 7 | 11 | 5 | 11 |
| Heart racing | 6 | 12 | 7 | 15 | 4 | 7 | 3 | 8 |
| Alert | 47 | 31 | 42 | 28 | 48 | 33 | 37 | 28 |

### Information Bias Learning Task

#### Model-Based Analysis

Losartan, relative to placebo, produced a significant positive versus negative bias in learning rates in the first block (valence x group interaction: F(1,49)=5.110, *p*=.028). Losartan suppressed loss learning rates relative to placebo (t(51)=-2.029, *p*=.048) while leaving win learning rates intact (t(51)=.562, *p*=.58). Win learning rates were significantly higher than loss learning rates in the losartan group (t(27)=3.301, p=.003), whereas these were comparable after placebo (t(24)=.032, p=.974) (Figure 2A).

**Figure 2.**
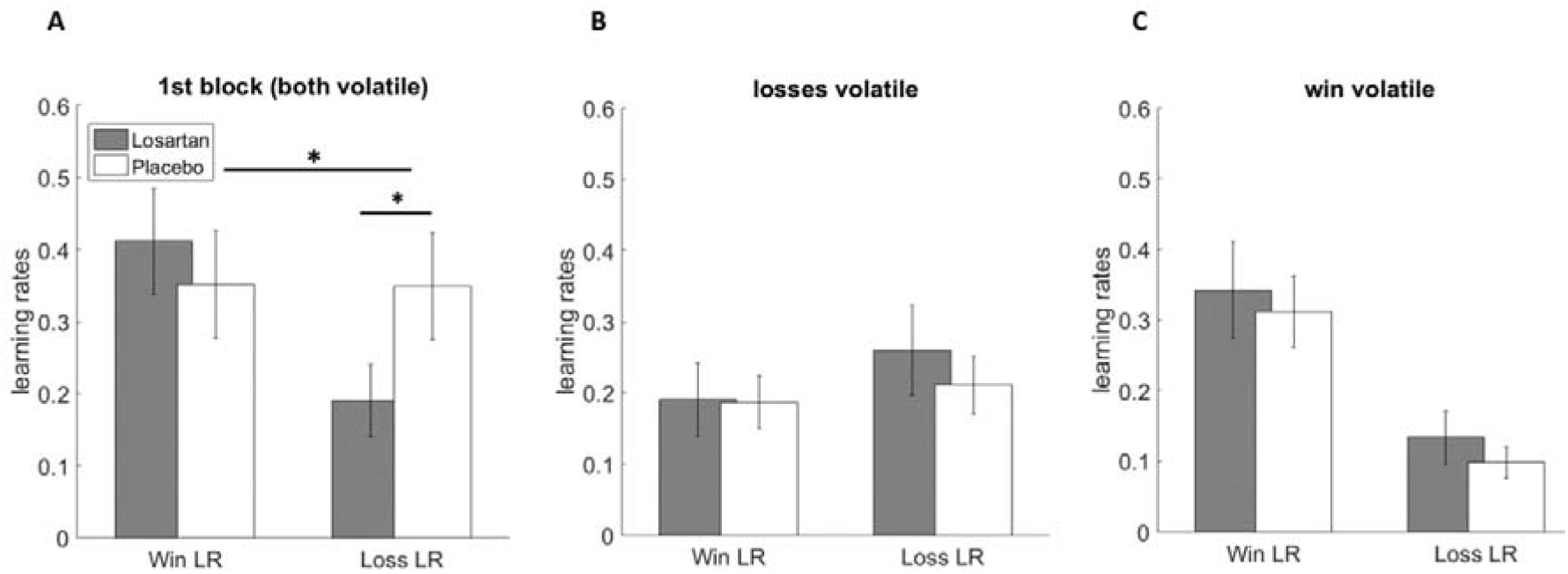
Behavioural effects of losartan on learning rates (LR) in different blocks of the IBLT. (A) Losartan had a specific effect on learning rates for losses relative to wins (F(1,49)=5.110, *p*=.028). More specifically, the losartan group had significantly lower loss learning rates than the placebo group (*p=.048). (B-C) Losartan did not alter learning rates in the later 2 blocks in which one outcome was volatile and the other stable. Learning rates for wins and losses are represented separately. The mean ± SEM are displayed.

Losartan did not influence learning rate adjustment in response to outcome volatility in the final two blocks (group x block x valence interaction: F(1,49)=.101, *p*=.752). Rather, all participants adjusted learning rates in response to outcome volatility as reported previously (29) (block x valence interaction: F(1,49)=41.942, *p*<.001) (Figures 2B, 2C).

The effect of losartan was specific to the learning rate parameter from the model, with no effect observed for either the inverse temperature or bias terms (Supplementary Figure 1, all F(1,49)<.902, all p>.407).

#### Non-Model-Based Analysis

Losartan increased the influence of win relative to loss outcomes on learning behaviour in the first block (group x trial type interaction: Wald’s *χ*^2^(1)= 4.87, *p*=0.027). Losartan participants switched choice significantly less often when selecting a shape associated with both win and loss (main effect group: Wald’s *χ*^2^(1)= 4.55, *p*=0.033), with the increase in switching when neither outcome was associated with the current choice being non-significant (main effect of group: Wald’s *χ*^2^(1)= 0.003, *p*=0.95). Consistent with model-based results, losartan had no effect on choice switching in the final two blocks (terms including group: Wald’s *χ*^2^(1)< 3.2, *p* > ^0^ .07). Losartan did not influence total amount of money won (t(51)=-.34, *p*=.73) or reaction time (F(1,49)=.06, *p*=.808) (Figure 3).

**Figure 3.**
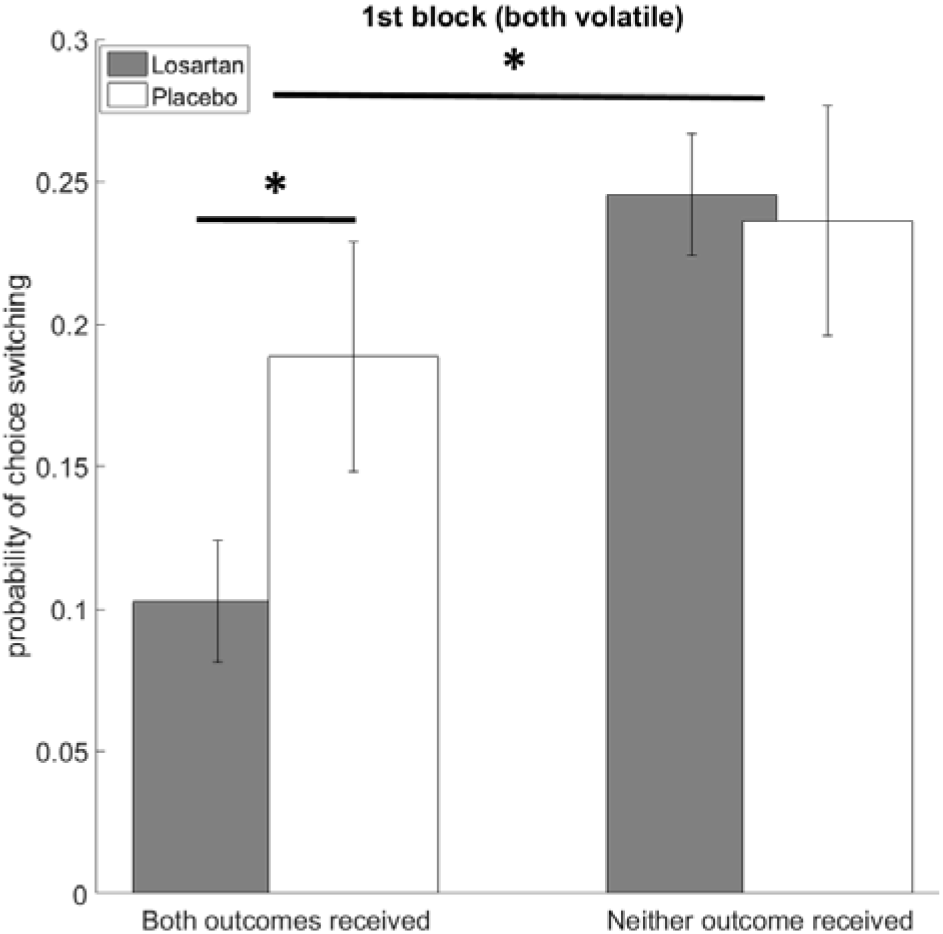
Choice switching analysis of learning behaviour in the IBLT. Losartan increased the influence of win relative to loss outcomes on learning behaviour in the first block of the IBLT task (group x trial type interaction: Wald’s χ^2^(1)= 4.87, *p*=0.027). Losartan significantly reduced participants’ probability of switching to the unchosen shape after receiving both win and loss outcomes in the first block of the task (main effect of group: Wald’s χ^2^(1)= 4.55, *p*=0.033).

## Discussion

We demonstrate that a single dose of the angiotensin II receptor antagonist losartan induces a positive learning bias in healthy human participants. More specifically, losartan reduced the degree to which participants were influenced by aversive outcomes, while leaving the influence of appetitive outcomes unaffected. These results suggest a potential cognitive mechanism by which losartan may influence the course of exposure therapy.

The task used in this study assessed the degree to which participant choice was influenced by both positive (wins) and negative (losses) outcomes. This was possible because the outcomes were independent, in other words knowing where the win outcome would occur provided no information about where the loss outcome would occur and vice versa. As a result, participants had to separately learn the likely location of the two outcomes, allowing the influence of these estimates on choice behaviour to be assessed. Losartan exerted a clear effect on choice switching behaviour in the first block of the task; the choices of participants who had received losartan were more influenced by win than by loss outcomes. Specifically, losartan participants were significantly more likely to select the same shape the trial after that shape had been associated with both outcomes and were less likely, albeit non-significantly, to switch their choice after choosing a shape associated with neither outcome. In other words, when the win and loss outcome drove choice behaviour in opposite directions, participants who had received losartan were more likely to make a choice prompted by the win outcome. While this analysis illustrates the basic effect of losartan, it only estimates the relative influence of the two outcomes and is therefore unable to determine whether participants were *more* influenced by the wins or *less* influenced by the losses. This level of analysis is possible using a computational modelling based approach in which learning rates for the two outcomes can be independently estimated. Taking this approach revealed that losartan exerted a specific effect, suppressing the degree to which participants learned from negative outcomes, while leaving learning from positive outcomes intact. In effect, participants who received losartan treated the loss outcome as being less informative than those who received placebo (23).

In contrast to its effect in the initial block of the IBLT task, losartan did not influence the degree to which participants adjusted their learning rates in response to outcome volatility in the last two blocks. The volatility of an outcome influences how informative that outcome is (23). Normatively, learning rates should increase as the volatility of the association being estimated increases, as this is the most efficient learning strategy in dynamic environments (37). That losartan did not alter this process indicates that participants were still able to identify which of the two outcomes was most informative during learning and tune their learning rates to this estimate. In other words, losartan caused participants to initially bias their learning away from aversive outcomes, but they remained able to learn from those aversive outcomes if they estimated them to be informative. Consistent with this, losartan did not impair the overall performance of participants in terms of total monetary wins, the inverse temperature parameter or reaction time.

A proposed key mechanism of action of exposure therapy is inhibitory learning, where the association of a stimulus with threat is suppressed by a new neutral or even positive association (5-7). A patient with panic disorder and agoraphobia might fear the onset of a heart attack if disconnected from the possibility of immediate medical help and therefore avoid situations such as large crowds or lonesome parks. Exposure tests out the anticipated aversive outcome of a heart attack and involves remaining in trigger situations for a prolonged period of time. The clinical effects of exposure are rooted in the patient making positive rather than the anticipated negative associations, leading to the anxiety decreasing gradually over time (38). In the present study, we found that losartan reduced learning from aversive events, while simultaneously preserving learning from positive outcomes. These effects are promising as they suggest that losartan might have the potential to also facilitate positive learning during exposure therapy in humans. Interestingly, we have recently reported that losartan increases the degree to which the amygdala differentially responds to appetitive and aversive stimuli in anxious participants (39). Better discrimination of appetitive and aversive stimuli, as demonstrated in this previous neuroimaging study, may represent an important mechanism by which losartan can induce the affective learning bias described in the current paper.

While the precise mechanism of action of losartan on learning and decision making is unclear, pre-clinical work provides some evidence that angiotensin II receptor activity may impact on dopaminergic function. Losartan binding to the angiotensin II receptor is associated with increased activity of the dopamine D1 receptor (albeit in kidney cells; (21)), while administration of losartan reduces the release of dopamine in response to nicotine in striatal preparations (20) and inhibits cell death in a dopaminergic cell line (22). Although this work provides no conclusive evidence for the role of angiotensin II receptors, it suggests that the effect of losartan on fear extinction in rodents (12) and affective learning as demonstrated by the current results may be mediated via an effect on dopaminergic transmission. More generally, this work is consistent with an impact of losartan on reward related processing, which has been linked to the development and treatment of anxiety disorders in humans. Research suggests that in anxiety disorders, neural response to reward is impaired (40, 41), and impaired reward response has been linked to future onset of anxiety and impaired response to exposure therapy (42, 43). In summary, while the detailed mechanism of action of losartan in exposure remains to be determined, the results from the current study are consistent with the idea that exposure-based treatments could be developed into more effective formats by combining them logically with pharmacological add-on compounds that synergistically target appetitive processing.

Similarly, our findings point to the possibility that losartan may prevent the development of anxiety symptoms which are triggered by unexpected traumatic events. A previous observational study has revealed that being treated for high blood pressure using angiotensin converting enzyme inhibitors or angiotensin receptor blockers such as losartan while experiencing a trauma is associated with a reduced tendency to develop traumatic symptoms, with no such effects were seen with other blood pressure medication (13). The results in the current study, where learning from aversive events was suppressed, may provide a cognitive mechanism for this observation. From a practical perspective, it would be interesting to test whether the effect of losartan on these posttraumatic symptoms is evident if the drug is initiated shortly after trauma, as this may suggest a use for the medication in prevention of PTSD.

While these results are promising for the development of more effective combination treatments for anxiety disorders, there are limitations to their interpretation. Even though our results provide evidence that losartan attenuates learning form aversive events, a mechanism proposed to be underlying successful exposure therapy (5-7), no assessment of the efficacy of losartan in exposure has been completed, and the moderating effect of the decrease in aversive learning following losartan on clinically meaningful outcomes of exposure therapy remains to be tested.

Taken together, this study has identified a specific effect of losartan on a computationally defined learning parameter and suggests a role for angiotensin receptors in learning about aversive outcomes. We provide evidence that a single dose of the drug, in the absence of overall effects on blood pressure, heart rate or transient mood changes, initiates a shift from aversive to appetitive learning. This mechanism has previously been suggested to be central to exposure therapy, hinting at the possibility that the effects observed here might positively interact with the clinical effects of exposure-based treatments. Such knowledge will facilitate the development of this and similar agents for optimal combination with exposure therapy, to maximally exploit synergistic effects.

## Acknowledgements

This research was funded by a MQ: Transforming Mental Health fellowship awarded to AR (MQ14F192).

## Supplementary Information

### Methods and Materials

In our previous work (23), we used a slightly different reinforcement learning model than the one reported in the main body of the article. The previous model included two inverse temperature terms and no bias parameter. The model reported in the main paper was selected as it provided a superior fit to participant choices (i.e. based on sum of Bayesian Information Criterion scores, 7676 versus 7604 where lower values indicate a better model fit). However, the learning rate effects reported in the current paper are not dependent on the specific model selected with the same pattern of significant results being found if the original model is used in the analysis.

For completeness, the values of the 4 free-parameters estimated under the best fitting model reported in the current manuscript, for the 1st block of the task in which we showed the effect of losartan, is visualised on a value map (Supplementary Figure 2).

### Results

**Supplementary Figure 1.**
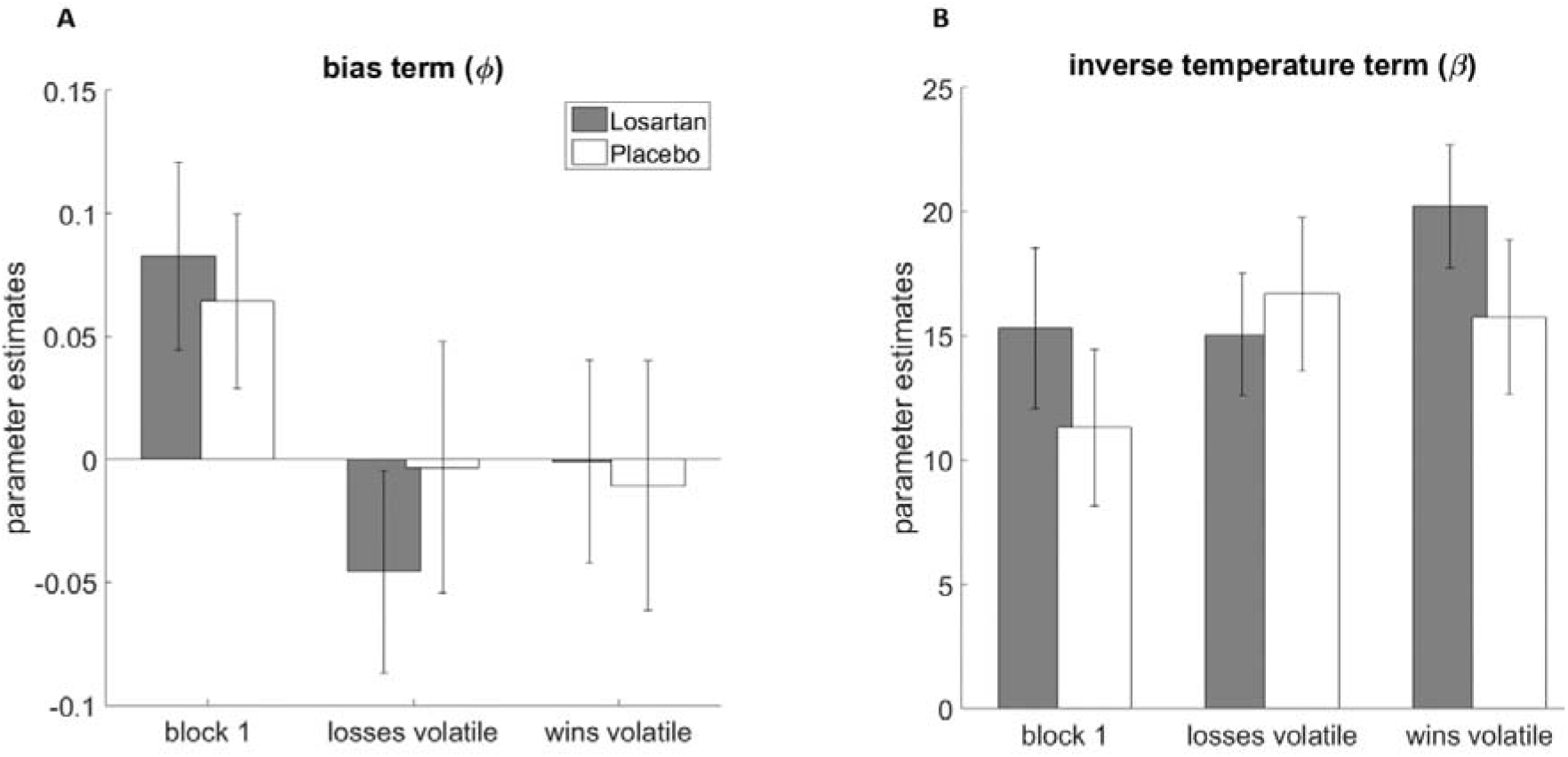
Effect of losartan on model parameters of no interest. **(A)** Administration of a single dose of losartan did not significantly influence other model parameters, such as the bias term accounting for unexplained biases to choose one shape over the other, or **(B)** the inverse temperature term accounting for participants’ choice stochasticity.

**Supplementary Figure 2.**
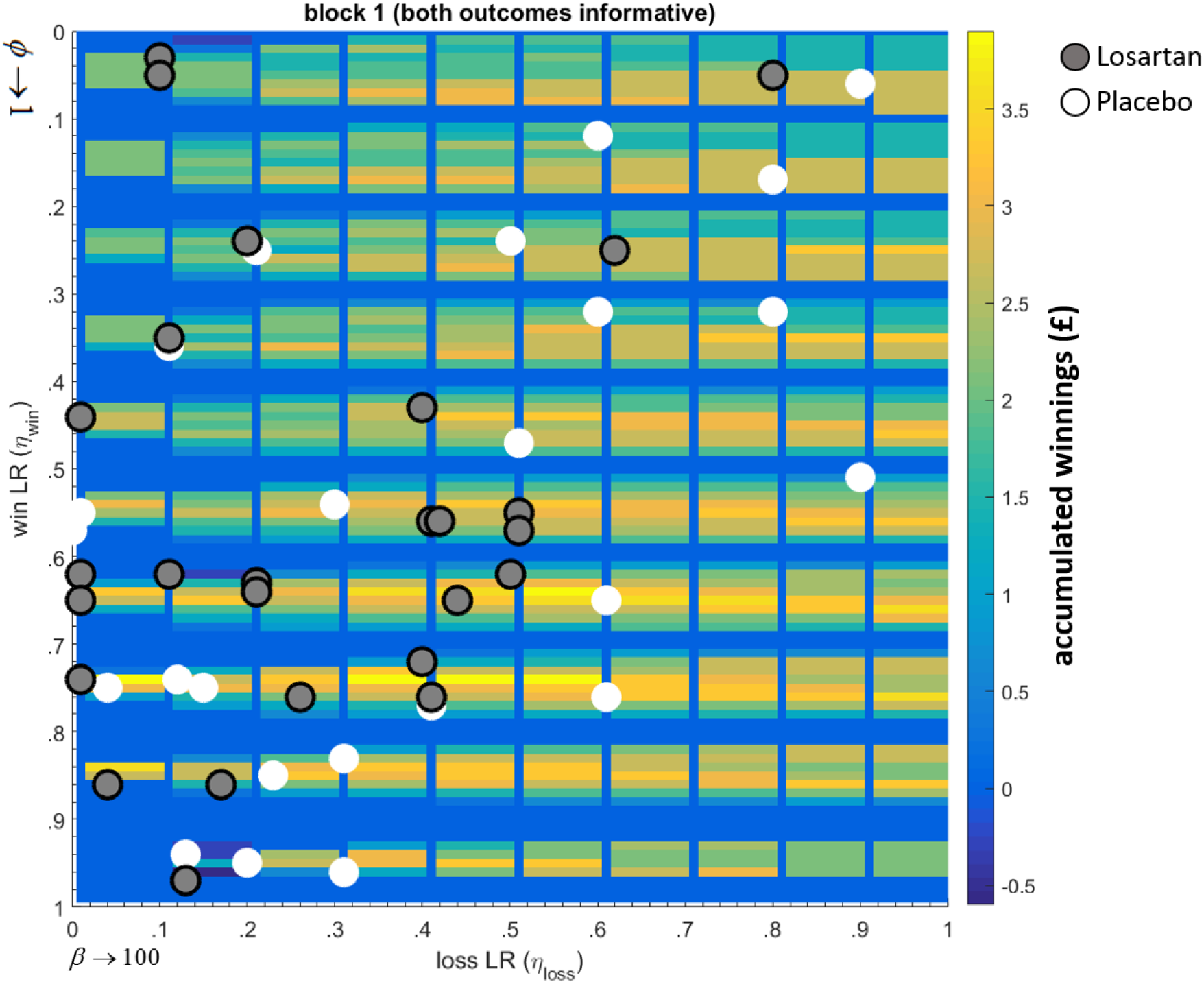
Overall behavioural effects of losartan expressed in terms of participant’s positioning on the Block 1 (both informative/volatile) value map. The best fitting model to participant choice behaviour had 4 free parameters, meaning that human behaviour can be expressed in terms of the combination of these parameters in an imaginary 4 dimensional space. For plotting purposes, we converted this parameter space to a 2D map where each axis defines win and loss learning rates. Within the y-axis and within each increment of the win learning rate, the bias term linearly increases from −1 to 1. Similarly, within the x-axis and within each increment of the loss learning rate, the inverse temperature term linearly increases from 0 to 100. A value map of block 1 is then constructed over the whole parameter space by generating responses by the best fitting model using the parameter combination at every cell of this grid, and calculating the estimated winnings based on these generated responses. Each participant’s estimated parameter values is then localised on this grid for a visual demonstration of overall how losartan influence human reinforcement learning. The figure demonstrates that the majority of participants in the losartan group fall into the left half of this value map defined by lower loss learning rates.

